# Stabilization of cultural innovations depends on population density: testing an epidemiological model of cultural evolution against a global dataset of rock art sites and climate-based estimates of ancient population densities

**DOI:** 10.1101/705137

**Authors:** Richard Walker, Anders Eriksson, Camille Ruiz, Taylor Howard Newton, Francesco Casalegno

## Abstract

Demographic models of human cultural evolution have high explanatory potential but weak empirical support. Here we use a global dataset of rock art sites and climate and genetics-based estimates of ancient population densities to test a new model based on epidemiological principles. The model focuses on the process whereby a cultural innovation becomes endemic in a population. It predicts that this cannot occur unless population density exceeds a critical value. Analysis of the data, using a Bayesian statistical framework, shows that the model has stronger empirical support than a null model, where rock art detection rates and population density are independent, or a proportional model where detection is directly proportional to population density. Comparisons between results for different geographical areas and periods yield qualitatively similar results, supporting the robustness of the model. Re-analysis of the rock art data, using a second set of independent population estimates, yields similar results. We conclude that population density above a critical threshold is a necessary condition for the maintenance of rock art as a stable part of a population’s cultural repertoire. Methods similar to those described can be used to test the model for other classes of archaeological artifact and to compare it against other models.

## Introduction

It is widely accepted that the complexity and diversity of human cultures are a result of Cumulative Cultural Evolution (CCE), enabled by humans’ unique neuroanatomy and cognitive capabilities, especially their skills in “cultural learning” [1–3]. However, the old idea that modern human capabilities emerged suddenly as a result of an advantageous mutation some 50,000 years ago [4,5] is no longer accepted and many allegedly modern features of human behavior and cognition are now believed to be more ancient than previously suspected [6–9]. There is, furthermore, no evidence that variations in brain size, brain morphology or innate capabilities explain any aspect of the spatiotemporal patterning of human cultural evolution over the last 50,000 years. Against this background, demographic models of CCE [10] assign a determining role to variations in population size, density and structure. Such models suggest that larger, denser, better connected populations produce more, more complex innovations than smaller ones [11], are less vulnerable to stochastic loss of unique skills [12,13], and are more capable of exploiting beneficial transmission errors during social learning [14]. They also suggest that well-connected metapopulations produce faster cultural innovation than metapopulations with weaker connections among subpopulations [15].

Demographic models could potentially provide valuable explanations for spatiotemporal patterning in the global archaeological record and several authors have used them for this purpose e.g. [14,16]. However, the assumptions and conclusions of the models are hotly contested [17–19]. Empirical studies are sparse and inconclusive, with some supporting an important role for demography (e.g. [14,15,20,21]), while others find no evidence for such a role [22–26].

In general, previous empirical studies have focused on the complexity of the broad classes of technology (e.g. food gathering technology) available to historical or recently extinct hunter-gatherer populations from specific geographical regions. Here, by contrast, we test the ability of a model to predict the global, spatiotemporal distribution of a single class of artifact (parietal rock art) over a period of 46,300 years.

## Results and discussion

### The model

CCE involves the inception, diffusion and selective retention of cultural innovations (e.g. innovations in social practices, beliefs, and technologies). In the process, a subset of the innovations become *endemic –* providing a stable foundation for future innovation.

In this study, we consider the case of parietal rock art. The model tested here (see Fig 1), inspired by SIR models in epidemiology [27–29], focuses on the process whereby an innovation becomes a permanent feature of a community’s cultural repertoire. We treat this process as analogous to the way a disease becomes endemic in a population [30].

**Fig 1.**
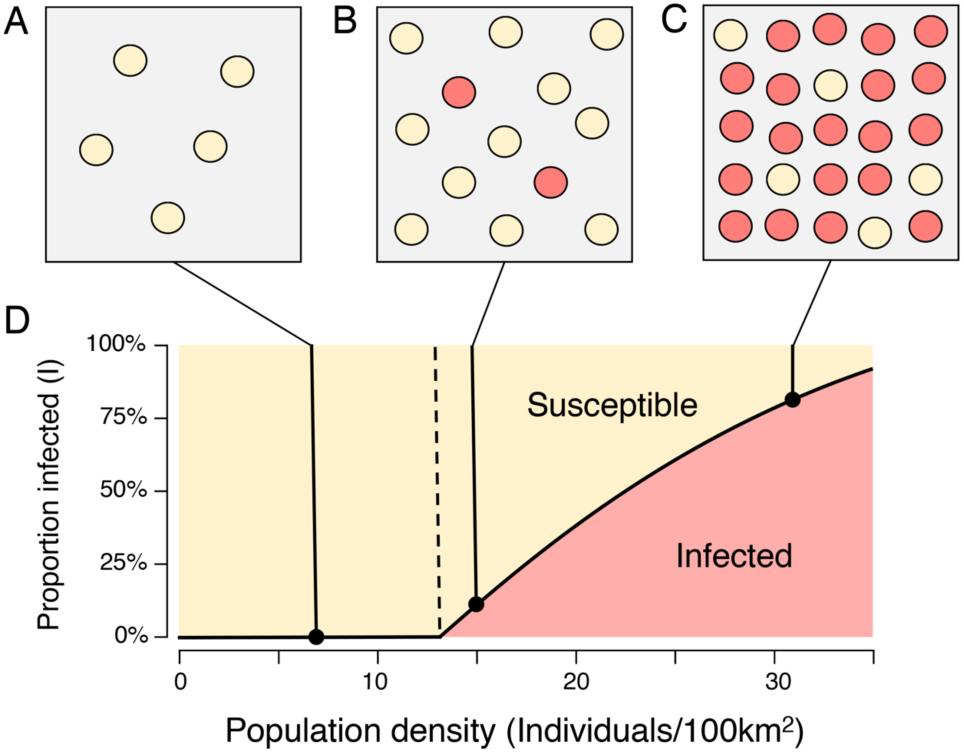
The epidemiological model. As population density increases, opportunities for transmission between subpopulations also increase. A: Below the critical threshold ρ^*^, the innovation is rapidly extinguished. B-C: Above the critical threshold, the proportion of infected subpopulations is an increasing function of population density.

Consider the emergence of rock art in a metapopulation comprising N subpopulations or communities, where I and S are respectively the proportion of infected communities (communities that have adopted the innovation), and susceptible communities (communities that have not). The innovation is transmitted from infected to susceptible communities at rate *β*. By analogy with empirical findings in [31] (see Methods), rates of encounter between communities are assumed proportional to the square root of population density. Infected communities recover (here: lose the ability or the propensity to produce rock art) at rate γ. As envisaged in [12,13], small communities, where only a few individuals possess a given skill, are particularly vulnerable to such losses.

In this model (see Fig 1 and Methods), there is a critical population density:

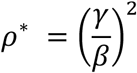

such that

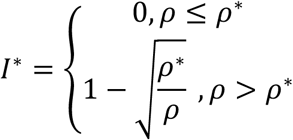

The value of *ρ*^*^ is determined by the equilibrium between the rate of transmission, *β*, and the rate of recovery, γ. Below *ρ*^*^, no communities are permanently infected. Above *ρ*^*^, the innovation spreads from community to community until the number of infected communities is equal to *I*^*^: in other words, it becomes *endemic*: even when a particular subpopulation loses the innovation it can reacquire it from other subpopulations where it has been retained. Examples of the loss and subsequent reacquisition of lost technologies are well-attested in the anthropological literature (see for example the reported loss and reintroduction of kayak-building skills among the Iniguit people [32,33]). The analogy with endemic disease is clear.

*I*^*^ is not directly observable in the archaeological record. To test our model, we therefore consider the *site detection rate* i.e. the probability, P, that a single cell in a hexagonal lattice of equally sized cells covering the surface of the globe, contains at least one recorded instance of rock art. Since the proportion of cells where geology, climate and research effort allow the creation, preservation and recording of rock art is small,

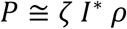

where *ζ* reflects the joint effects of these factors (see SI). The model predicts that for *ρ* ≤ *ρ*^*^, detection rates will be zero and that at higher densities it will rise approximately in direct proportion to *ρ*. As a result, the distribution of population densities for cells containing rock art will differ significantly from the distribution for all cells. Similarly, the median population density for cells containing sites will be automatically higher than for the complete set of cells.

### Testing the model

#### Assumptions and strategy

Our analysis is based on the assumption that an innovation that does not become endemic is statistically unlikely to contribute to CCE or to leave a trace in the archaeological record. If this is correct, most known rock art will originate in locations and periods where rock art was once endemic. The model predicts that all such sites will be located in territories whose population density exceeds a critical threshold.

To test these predictions, we collated a dataset containing the locations, dates and other characteristics of 133 scientifically dated rock art sites (see Fig 2A, SI Table 1, Methods). Uncalibrated radiocarbon dates in the dataset were calibrated using Calib 7.0 [34] with the IntCal 13 calibration curve. Fig 2B shows the resulting age distribution. Ancient population densities were estimated by combining data from two published models, informed by global genetic variation, and climate-based estimates of Net Primary Productivity [35,36] (See Methods). Using the results from these models, and applying the same methods, we represented the distribution of population densities in time and space on a lattice of *cells*, each approximately 100 km wide, and each associated with a date. Cells containing at least one example of dated rock art were defined as *sites*. In the subsequent analysis, we filtered this initial data set to include only cells lying at latitudes between 20-60°N and 10-40°S, and with dates more recent than 46,300 years ago - the only latitudes and dates with significant numbers of sites in our rock art dataset. Cells with inferred population densities of zero were also eliminated. This process led to the exclusion of 4 sites at equatorial latitudes, 1 site older than 46,300 years and 8 sites with inferred population densities of zero. After filtering, the final dataset included 9.8 million cells containing 119 rock art sites.

**Table 1:** Summary of statistical results for the analyses described in the text. The Kolmogorov-Smirnov test tests the null hypothesis that the distribution of population densities for *sites* does not differ significantly from the distribution for all cells. The Mann-Whitney test (which is less sensitive) tests for differences between the medians. Threshold CIs show the inferred confidence intervals for the threshold, given the empirical data. Log likelihood shows the log likelihood of the model, given the empirical data. The two Bayes factors show the ratio of the marginal likelihood of the epidemiological model to the marginal likelihood of alternative models (see Methods)

| Analysis | Eriksson | Eriksson exact direct dates (calibrated by original authors) | Timmermann | Australia | France-Spain | 0 – 9,999 years ago | 10,000-46,300 years ago |
| --- | --- | --- | --- | --- | --- | --- | --- |
| N sites | 119 | 37 | 93 | 56 | 31 | 74 | 45 |
| N. cells | 9,817,159 | 9,817,159 | 213,207 | 1,223,500 | 242,009 | 2,672,852 | 7,144,307 |
| Median population density sites | 29.77 | 30.30 | 25.04 | 30.28 | 28.23 | 30.34 | 27.04 |
| Median population density all cells | 15.29 | 15.29 | 18.04 | 21.50 | 26.60 | 19.92 | 13.65 |
| KS D | 0.58 | 0.73 | 0.54 | 0.64 | 0.68 | 0.76 | 0.45 |
| KS p | 0.000016 | 0.000000 | 0.000336 | 0.000001 | 0.000000 | 0.000000 | 0.000132 |
| Mann-Whitney U | 327.00 | 241.00 | 236.00 | 245.00 | 261.00 | 205.00 | 403.00 |
| Mann-Whitney p | 0.002600 | 0.000034 | 0.005337 | 0.000048 | 0.000830 | 0.000005 | 0.034209 |
| Threshold CI 0.025 | 7.36 | 9.64 | 12.80 | 11.84 | 0.11 | 8.96 | 0.92 |
| Threshold CI 0.25 | 10.72 | 13.76 | 15.64 | 17.30 | 6.23 | 13.18 | 4.26 |
| Threshold CI 0.5 | 12.27 | 15.41 | 17.19 | 26.81 | 14.12 | 15.03 | 6.83 |
| Threshold CI 0.75 | 14.05 | 16.77 | 18.94 | 28.00 | 18.56 | 16.53 | 9.53 |
| Threshold CI 0.975 | 16.64 | 18.65 | 21.79 | 28.90 | 32.83 | 18.82 | 14.53 |
| Log Likelihood | -1404 | -467 | -791 | -593 | -303 | -820 | -560 |
| Bayes Factor (w.r.t. proportional model) | 1.07E+05 | 1.54E+05 | 2.55E+03 | 1.47E+04 | 2.00E+01 | 3.88E+03 | 3.49E+01 |
| Bayes Factor (w.r.t. Constant model) | 9.79E+26 | 6.33E+13 | 2.22E+09 | 8.72E+09 | 2.47E+02 | 2.29E+13 | 1.50E+10 |

**Table 2:** Definition of “All cells” used in the analyses

| Analysis | Definition of “all cells” |
| --- | --- |
| World | All cells with non-zero population with dates in the range 0-46,300 years ago and locations between 60°N and 20°N or between 10°S and 40°S |
| Periods | (1) All cells with non-zero population with age between 0 and 9,999 years.<br>(2) All cells with non-zero population with age between 10,000 and 46,300 years. |
| France and Spain | Derived from the maximum and minimum latitudes and longitudes for sites in our dataset with locations in modern France or Spain. All cells with non-zero population lying in a “rectangle” with NW corner (lat: 50.84°N, lon: 10°) and SE corner (lat: 35.00°N, lon 7.00°E) |
| Australia | Derived from the maximum and minimum latitudes and longitudes for sites in our dataset with locations in modern Australia. All cells with non-zero population lying in a “rectangle” with NW corner (lat: 25.27°S, lon: 133.77°E) and SE corner (lat: 39.16°S, lon: 154.86E) |

**Fig 2.**
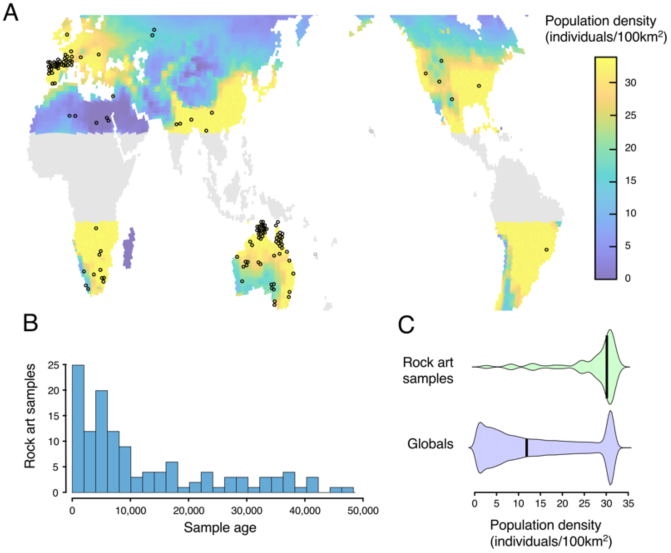
Sites used in the analysis: A: Geographical distribution of sites (all 133 sites) and inferred population distributions (maximum value over the last 46,300 years, see color bar for scale; areas excluded from the analysis shown in grey). B: Distribution by earliest date of rock art at site location (119 sites included in analysis). C: Comparison of population densities for sites vs. all cells (119 sites included in analysis).

Using this dataset, we computed the distribution of population densities for sites and for all cells (Figure 2C). On this basis, we calculated site detection rates (number of sites / total number of cells) for each population density bin.

### Empirical support for the model

#### Full dataset

As a first test of the model, we analyzed population densities for our complete rock art dataset (119 sites). As predicted, the distribution of population densities for sites differed significantly from the distribution for all cells (two sample KS test D=0.58, p<0.0001), the proportion of sites in cells with low population densities was much lower than the proportion of all cells, and the proportion in cells with high densities was much higher (median population densities: Sites: 29.77/100km^2^; All cells 15.29/100km^2^, Mann-Whitney U=327, p<0.01) (See Table 1 for detailed results).

Using a Bayesian statistical framework, we estimated the most likely parameter values for the model given the empirical observations (Fig 3A). Posterior distributions for model parameters were tightly constrained (see SI Fig 1). The inferred critical population density for the model with the highest likelihood, was 12.27 individuals/100km^2^ (95% CI: 7.36 – 16.64) (Fig 3B). Comparison of the model against a constant model (equal site frequency at all population densities) strongly supported rejection of the alternative model. Comparison against a proportional model, where detection rates were directly proportional to population density (i.e. to the number of potential inventors in the population) (compare with [11]), again supported the superiority of the epidemiological model. (for detailed results see Table 1). Taken together, these findings support the hypothesis that endemic production of rock art requires population density above a critical value.

**Fig 3.**
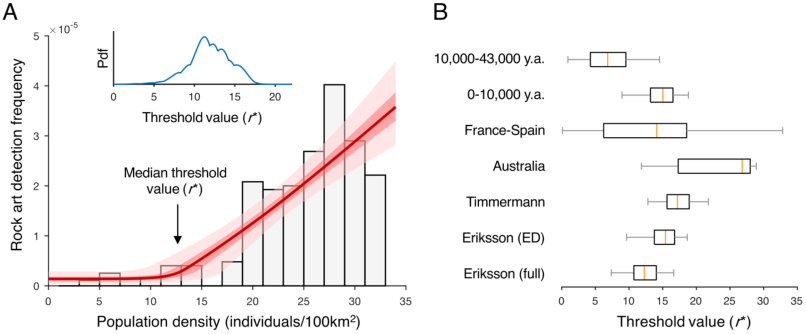
Empirical support for the epidemiological model. A. Inferred detection frequencies given the epidemiological model, the full archaeological dataset, and the combined population estimates from [35,36] (see Methods). Grey bars: Empirical frequencies of rock art in different intervals of population density. Red line: Estimated rock art detection rate as a function of population density (median of posterior distribution of estimated detection rate, with interquartile range in dark pink and 95% CI in pale pink). Also shown is the posterior probability density function (pdf) of the threshold value (ρ^*^, inset); the median value of this distribution is indicated by an arrow on the main axis. B. Inferred values of the critical threshold for the different analyses described in the paper (ED: exact direct; orange line: maximum likelihood estimate; box: interquartile range; whiskers: CI 0.025-0.975).

### Sensitivity to potential dating errors

Eight sites in the rock art dataset were located in cells with estimated population densities below the critical value inferred from the model and the empirical data. As described in the Methods section, some of the dates in our rock art dataset were minimum or maximum dates, and some were estimated using indirect methods (e.g. dating of organic materials stratigraphically associated with the art). Additionally, some radiocarbon dates (mainly from the older literature) were not calibrated or came from papers that did not report their calibration status. Since these dates do not necessarily reflect the actual dates at which the art appeared, and correct inference of population densities requires correct dates, we repeated our analysis using only sites with “high quality” dates, i.e. exact dates obtained with direct methods, and, in the case of radiocarbon dates, dates calibrated by the original authors.

In this filtered dataset, all sites (37/37) had population densities higher than the inferred critical population density (see below). This finding matches the predictions of the model and suggests that the unexpectedly high inferred population densities for some of the sites in the original dataset, may have been due to erroneous dating. As in the original analysis, the distributions of population densities for sites differed significantly from the distribution for all cells (Two sample KS test D=0.73, p<0.00001), median population density for sites was higher than for all cells (Sites: 30.30; All cells 15.29, Mann-Whitney U=241, p<0.0001) and, empirical support for the epidemiological model was much stronger than for alternative models (see Table 1). The 95% CI for the inferred critical population density (15.41 individuals/100km^2^, 95% CI: 9.64-18.65) overlapped with the estimate for the complete dataset (see Figure 3B, Table 1).

#### Alternative population estimates

Another potential source of errors was the model used to generate our population estimates. We therefore repeated our analyses using more recent population estimates [37], based on a different climate model and different assumptions (see Methods). As in the earlier analysis, there were significant differences between the population density distributions for sites and all cells, (two sample KS-test D=0.54, p<0.001), median inferred population was higher for sites than for all cells (sites: 25.04; all cells: 18.04, Mann-Whitney U=236, p<0.01) and empirical support for the epidemiological model was much stronger than for the alternative null or proportional models (see Table 1). The CI for the inferred critical population density (95% CI: 12.8-21.79) overlapped with the CI for the original analysis (see Fig 3B, Table 1)

#### Potential sample bias

A comprehensive survey of the vast rock art literature was beyond the scope of our study. As a result, large geographical areas in Central/South America, Central Africa, East and South-East Asia are practically unrepresented in our dataset (see Fig 2A). Moreover, 60.1% of the sites in our full dataset date from less than 10,000 years ago, and only 3.9% from before 40,000 years ago (see Fig 2C). All this suggests the possibility of systematic bias. Suggestions that the literature itself is biased by ecological and taphonomic factors and by geographical variations in research effort [38] make the presence of such biases even more plausible.

To test the potential impact of such biases, we repeated our analysis for data from two culturally unrelated geographical areas (the territory covered by the modern countries of France and Spain, and Australia) and for two distinct periods (from the present until 10,000 years ago, and from 10,000 until 46,300 years ago (see Methods). In the case of France-Spain, it was not possible to test the prediction of a critical threshold because nearly all cells had relatively high inferred population densities. The other three analyses supported the existence of such a threshold. All four analyses found different density distributions for sites and for all cells, higher median population densities for sites than for all cells, and stronger support for the epidemiological model than for alternative models (Fig 3B, Table 1). The convergence of results for different periods and geographical areas is evidence that the overall results of our analysis are not due to sampling biases (differences in sampling rates between periods or areas).

#### Population density as a proxy for social contacts

Another potential problem was the use of population density as a proxy for contacts between subpopulations. Grove suggests that individual encounter rates depend not on population density but on the product of density and mobility [31]. However empirical results included in the same paper show that, in reality, the mobility of modern hunter gatherers is inversely proportional to the square root of population density. If this is correct, the encounter-rate will be directly proportional to the square root of population density. This is the assumption on which we based our model. Testing with a generalized model (results not shown) demonstrated the robustness of our qualitative findings to the exact specification of this relationship (Methods).

## Conclusions

This study models one of the key mechanisms required for CCE (the process whereby an innovation becomes endemic in a population), and tests the predictions of the model for the case of parietal rock art. In all our analyses, (Fig 3B, Table 1), the distribution of population densities for sites differs significantly from the distribution for all cells and median population densities for sites (25-30 individuals/km^2^) are consistently higher. In all cases, empirical support for the epidemiological model and a critical population density is much stronger than for alternative proportional or null models. Taken together, these results provide robust grounds to reject the null hypothesis that the emergence of rock art is unrelated to demography, and strong support for our model i.e. for the hypothesis that population density above a critical value is a *necessary* condition for an innovation to become endemic in a population.

Importantly, nothing in our model or empirical results suggests that a minimum level of population density is a *sufficient* condition for the emergence of rock art. Our model represents a single aspect of Cultural Evolution – namely, the process whereby an innovation becomes endemic in a population. We thus consider our model as complementary to the demographic models of the social transmission and selective retention of cultural innovations, reviewed in the Introduction. It is certain, furthermore, that rates of preservation and detection of rock art are strongly dependent on climatic and geological factors, and on variations in research effort. It is not surprising, therefore, that many areas of the globe (e.g. in equatorial Africa) with high inferred population densities for the relevant periods, have little or no reported rock art.

Previous demographic models have posited a relationship between cultural evolution and the overall size of so-called Culturally Effective Populations, networks of cultural exchange spanning potentially large geographical areas. The theoretical arguments for this relationship are strong. However, in archaeological contexts where a local population’s cultural links to other populations are unknown, the size of “culturally effective populations” is hard to estimate [12]. One of the key methodological features of our study is the use of population density as the independent variable. Since population densities are easier to estimate than culturally effective population sizes, this greatly facilitates the process of theory testing. A second key feature is the choice of “detection frequency” as the dependent variable. In the majority of current studies the dependent variable is “cultural complexity”, usually represented by the size of the population’s toolkit [39]. The method used here circumvents the difficulty of applying such a concept in archaeological settings, where complete toolkits are rarely available. These methodological features of our study will facilitate the application of our model and methods outside the case of rock art and in contexts where other constructs are difficult to measure.

Our rock art dataset, population estimates, and software tools are publicly available at https://github.com/rwalker1501/cultural-epidemiology.git. Other researchers are encouraged to use them to replicate our findings, to test our hypotheses for other classes of artifact and with other population estimates, and to explore their own models.

## Methods

### Derivation of the model

The model describes the diffusion of a cultural innovation through a Culturally Effective Population (CEP) [12] i.e. a closed network of communicating individuals or subpopulations. Notation for the “disease dynamics” follows the conventions established in [29].

Consider a CEP with stable number of subpopulations, N. Assume that the population is divided into infected and susceptible (unaffected) subpopulations, with proportions *I* and *S*, respectively, such that *I* + *S* = 1. If the population is “well-mixed” (i.e. every subpopulation is in social contact with every other subpopulation), any infected subpopulation can transmit the innovation to any susceptible subpopulation. Let *β* be the rate of transmission per time step and γ the rate of recovery of the infected subpopulation (here: the rate at which it loses the ability to produce rock art). Thus, the rate of change in the proportion of infected subpopulations is given by

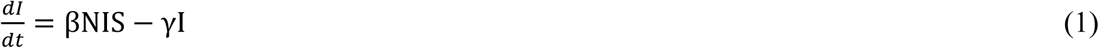

In a partially connected population, infected subpopulations can only transmit an innovation to the susceptible subpopulations with which it is in contact. Assuming that on average each subpopulation is in contact with k other sub-populations, we can substitute *k* for *N* in Eq. (1). Using *S*+*I*=1, we thus obtain

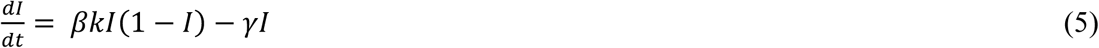

Rearranging, solving for 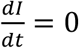, and imposing *I*^*^ ≥ 0, we obtain the stable proportion of infected subpopulations:

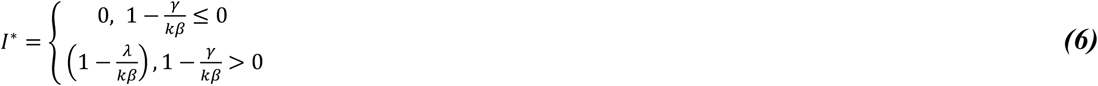

If social contact numbers k are proportional to the square root of density [31], i.e. 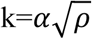,

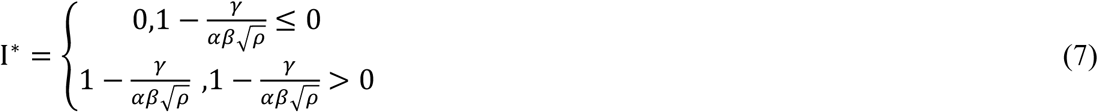

In short, there exists a critical population density, ρ^*^, such that at densities below ρ^*^ there are no infected subpopulations, and at densities above ρ^*^, the number of infected subpopulations is greater than zero.

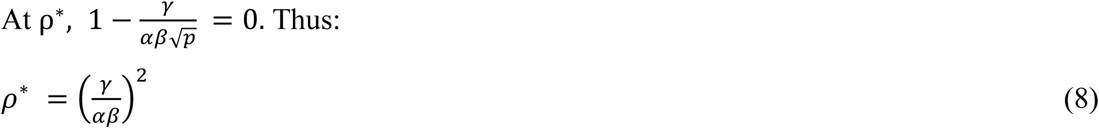

To reduce the number of parameters in the model, we normalize the rate of transmission, setting *αβ* to 1. We thus obtain the simplified expressions:

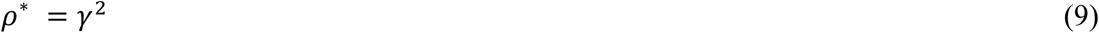

and

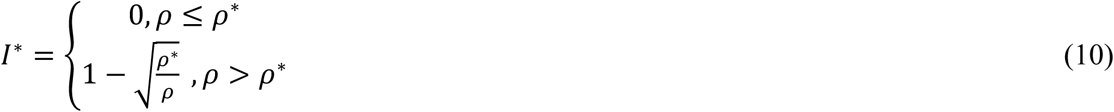

In this simplified version of the model, the equilibrium size of the infected population and the critical value of *ρ* are entirely dependent on population density and a single parameter, γ.

Finally, we assume that each infected subpopulation has an equal probability, *z*, of producing an artifact that produces a trace in the archaeological record. On this assumption, the probability *P*, that the n infected subpopulations will generate at least one such trace, is given by:

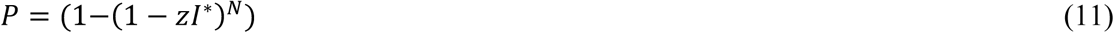

Assuming z is small,

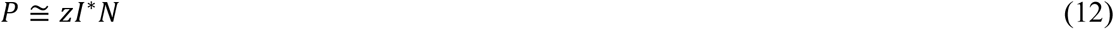

The value of N for a cell of standard area, with fixed mean community size, is proportional to the population density for the cell:

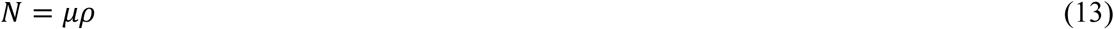

Grouping *μ* and z in a single variable *ζ* we obtain

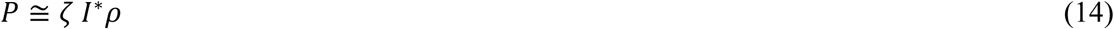

In the setting of our empirical study, inferred population sizes may contain large errors, especially when site dates are minimum or maximum dates or indirect estimates or when the calibration status of a radiocarbon date is not clearly stated in the original article. We, therefore, include an error term *ε*, representing the probability that a site is attributed to a cell with an incorrect date, whose population will thus be different from that of the cell with the correct date. Thus:

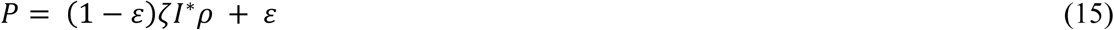

This is the model we tested in our empirical study.

### Rock art dataset

The analysis presented here is based on a dataset of parietal rock art, generated through a literature search with Google Scholar. We are aware that the database contains only a small proportion of all rock painting sites in the world, and that it may be subject to systematic biases. These are discussed in the Results and Discussion.

For the purposes of our survey, parietal rock art was defined to include all figurative and non-figurative pictograms (paintings and drawings) and petroglyphs (engravings) using rock wall as a canvas. “Portable art” (e.g. figurines) and geoglyphs (i.e. large designs or motifs on the ground) were excluded from the analysis.

The survey was seeded using the query:

(“rock art” OR “parietal art” OR petroglyphs OR pictographs) [AND] (radiocarbon OR AMS OR luminescence OR Uranium).

We read the top 300 articles found by the query that were accessible through the EPFL online library, together with other relevant papers, cited within these articles. Sites where drawings, paintings and engravings were reported, were systematically recorded. Sites with no radiocarbon, optical luminescence or Uranium-Thorium date were excluded.

For each dated site, we recorded the longitude and latitude of the site (where reported), its name, the earliest and latest dates of “art” found at the site (converted to years before 1950). Where authors reported a confidence interval for dates we used the midpoint of the confidence interval. Radiocarbon dates marked by the original authors as calibrated dates were flagged as such. The calibration status of radiocarbon dates whose calibration status was unspecified was inferred from the surrounding text and from the CIs for the dates. Dates inferred to be uncalibrated were then calibrated using Calib 7.0 [34] with the IntCal 13 calibration curve. Where different authors reported different dates for a site, without disputing dates proposed by other authors, we systematically used the dates from the most recent report. We also recorded the name of the modern country where the site was located, the journal reference, the method(s) used to produce the date, the nature of the dating procedure (direct dating, indirect dating), the nature of the data provided (exact data, minimum date, maximum date, mixed), a descriptor of the artifacts found (paintings, drawings, petroglyphs etc.), and a flag showing disputed dates.

In cases where the article did not provide a latitude and longitude, online resources were used to locate the information. The main such resources were D. Zwiefelhofer’s FindLatitudeAndLongitude web site [40], which is based on Google Maps, and Austarch, a database of 14C and Luminescence Ages from Archaeological Sites in Australia, managed by A. Williams and Sean Ulm [41].

The survey generated 190 records. Records with identical latitudes and longitudes and overlapping dates were merged (5 records eliminated). Duplicate records (12), modern sites (1), sites which did not meet the inclusion criteria (13), sites where the source was deemed unreliable (5), sites where reliable geographical coordinates were unavailable (12), sites with disputed or doubtful dates (8) and 1 site described in a retracted article were excluded. These procedures left a total of 133 records.

All except 5 of these records referred to sites located between 20°-60°N and 10° - 40°S and with dates more recent than 46,300 years ago. The most relevant comparison was thus with all cells *in the same range of latitudes*. Including a small number of sites at latitudes north or south of this range would have required a vast expansion of this definition, and the inclusion of many cells with extremely low population densities. Sites located outside this range of latitudes were therefore excluded from the subsequent analysis. For similar reasons, one very early site (Blombos Cave, South Africa) (77,000 years ago) was also excluded. After these exclusions, the dataset comprised 127 records. Subsequent analysis (see below) showed that 8 records referred to sites with inferred population densities less than 1 individual / 100km^2^ at the date corresponding to the earliest rock art at the site. These sites were also excluded. Thus, the final analysis was based on a dataset of 119 records.

### Estimates of population density

The population density estimates used in our paper combine results from a climate-informed spatial genetic model [35] with an improved model reported in [36]. Briefly, models combine climate estimates for the last 120,000 years, based on the Hadley Centre model HadCM3, with data on patterns of modern genetic variability from the human genome diversity panel-Centre d’Etude du Polymorphisme Humain (HGDP-CEPH), and a mathematical model of local population expansion and dispersal. Estimates of past precipitation and temperature are used to estimate Net Primary Productivity (NPP), and hence maximum human carrying capacity, for each cell in a hexagonal lattice with equal area cells 100 km wide, for all dates from 120,000 years ago to the present using time steps of 25 years. The carrying capacity is a continuous function of NPP governed by two NPP threshold values and a maximum carrying capacity parameter, K. The carrying capacity is zero below the lower NPP threshold, increases linearly from zero to K between the two NPP thresholds, and is constant and equal to K for NPP above the upper NPP threshold value. Human expansion out of Africa is simulated using a model where populations that have reached the maximum carrying capacity of a cell expand into neighboring cells. Approximate Bayesian Computing is used to compare models with a broad range of parameter values and starting values. Model evaluation is based on their ability to predict regional estimates of pairwise time to most recent common ancestor (TRMCA). Population density estimates for Eurorasia, Africa and Australia were computed using parameters from the high-ranking scenario described in Fig 2 and Movie S1 in [35]. The estimates for the Americas (considered as a single continent) were taken from [36]. Compared to the model presented in [35], this paper contains a more accurate model of ice sheet dynamics in the North of the American continent and related areas of Eastern Asia, more accurate estimates of NPP across the Americas and better dates for the interruption of the Beringian land bridge. Estimated dates for the colonization of the Americas, are more accurate than in the original model.

These population estimates were compared against the results using a second set of population estimates reported in [37]. The data refer to the early exit scenario (Scenario A) described in the paper. As in [35], human population density estimates are based on a climate model (LOVECLIM) combined with a reaction-diffusion Human Dispersal Model. Unlike the model in [35], the estimates do not take account of genetic data. A second difference concerns the population estimator, which is based not just on NPP but also on temperature and predicted “desert faction” and incorporates *ad hoc* modeling hypotheses absent in the previous model. These include accelerated human dispersal along coastlines (“a coastal express”) and a special Gaussian decay function, modeling the probability of island hopping as a function of distance. For clarity of presentation, we have converted population density estimates from the two models into the same units, namely effective population/ 100km^2^.

### Data analysis

In many sites in our dataset, the rock art reported in the literature spanned a broad range of dates. For the purposes of our analysis, we considered only the *oldest* date for each site. This procedure reduced our sample size and the statistical power of our analysis, but also the risk of artifacts due to dependencies in the data. Each site was associated with the estimated population density for the cell in the population model whose date and center point were closest to the date and location of the site. “All cells” were defined according to the needs of the individual analysis Data for sites and all cells were binned by population density (35 bins). Numbers of sites and all cells and frequency of sites (sites/all cells) were computed for each bin.

### Testing model predictions

To test the predictions of the model for a specific dataset, we compared the distribution of sites by population density against the equivalent distribution for all cells. Using the two-sided, two sample Kolmogorov-Smirnov test, we tested the null hypothesis that both were drawn from the same distribution. The difference between the medians of the two distributions was tested using the less sensitive Mann-Whitney U test.

Empirical support for the model was estimated using a Bayesian framework. For each of the parameters (γ, *ζ* and *ε*) we defined a uniformly distributed set of possible values lying within a plausible range and computed the likelihood of the model for all possible combination of these values, verifying the choice of parameter values using the computed posterior distributions. The value of the critical population density, *ρ*^*^ was inferred inserting the most likely inferred values of γ in the model

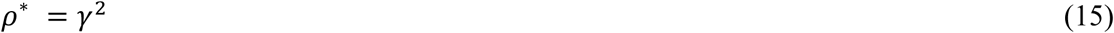

The most likely model was then compared against the most likely constant model, z(*ρ*) = *k*, and the most likely proportional model, z(*ρ*) = *ζρ*. The relative likelihoods of the model were quantified using Bayes factors. The results are included in Table 1.

### Sensitivity to potential dating errors

Some kinds of rock art (e.g. petroglyphs) make no use of organic pigments capable of providing an exact date for the artifact. In these cases, dating relies on the analysis of overlying and underlying materials providing minimum and/or maximum dates or on the analysis of other organic materials stratigraphically associated with the artifact. Dates obtained in this way are subject to large errors.

A further complication comes from the presence in our original dataset of studies with radiocarbon dates whose calibration status is not explicitly stated by the original authors and whose status is thus inherently ambiguous. It is possible, furthermore, that some uncalibrated dates, particularly from older studies, may themselves be of poor quality.

To test the impact of these potential sources of error, we repeated our analysis using only articles providing exact dates obtained directly from the artifact. In the case of radiocarbon dates, we restricted the analysis to sites with calibrated dates provided by the original authors.

### Analysis by period and geographical area

Subsets of “sites” and “all cells” belonging to specific date bands (0 – 9,999 years ago, 10,000 – 46,300 years ago) and specific geographical areas with high numbers of sites (France/Spain, Australia) were subjected to the same analyses applied to the full dataset.

### Sensitivity to the relationship between population density and encounter rate

To test the sensitivity of our model to the assumed inverse quadratic relationship between population density and encounter rate, we formulated a generalized version of our original model, where the critical population density and the encounter rates are power functions of population density. Thus:

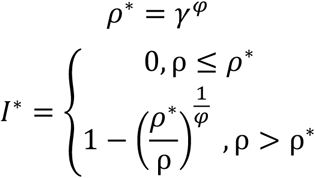

Replication of our analysis using this generalized model (results not shown) showed that our qualitative empirical findings are robust to changes in the specification of *φ*.

### Software

Data extraction and analysis was based on custom code written in Python 2.7, using the Anaconda development environment.

#### Software availability

Python source code for the software used to perform the analyses is available under a GPL 3.0 license, at https://github.com/rwalker1501/cultural-epidemiology.git. The software includes (i) the script used to generate the figures and tables shown in this paper; (ii) methods to run additional data analyses and to produce figures not shown in the paper; (iii) a menu driven program providing easy to use access to these functions; (iv) documented source code for the statistical calculations and plots used in the paper and the additional analyses

### Data

The GIT repository includes the full rock art database used for the analysis, and copies of the population estimates from the Timmermann and the Eriksson models (reproduced with the permission of the authors). The original population estimates from Eriksson can be found at: https://osf.io/dafr2/ The original population estimates from Timmermann can be found at https://climatedata.ibs.re.kr/grav/data/human-dispersal-simulation

## Supporting information

SM Table 1 (CSV)

## Acknowledgements

The idea of testing our model on the case of parietal rock art emerged from a meeting at University College, London, between Richard Walker, Stephen Shennan & Mark Thomas (University College, London) and Anders Eriksson. Axel Timmermann, International Pacific Research Center, University of Hawaii kindly contributed population estimates. Michelle Langley, Griffith University, Brisbane, Australia; Australian National University, Canberra, Australia, contributed data from a published survey of Australian rock art. Werner van Geit, Blue Brain Project, EPFL, Switzerland, contributed important code fragments. Isabel Marquez cross-checked the data for rock art sites. Mark Thomas and Stephan Shennan, Michael Herzog (EPFL Switzerland), Ruedi Füschlin (ZHAW, Switzerland), Lenwood Heath (Virginia Tech, USA) and Francesco Walker (University of Enschede, the Netherlands) reviewed earlier versions of this manuscript, providing valuable feedback. Henry Markram, leader of the Blue Brain Project, provided vital encouragement and support.

## Author contributions

RW: conceptualization, methodology, data curation, formal analysis, software, writing

AE: conceptualization, methodology, formal analysis, software, writing

CR: software

TH: formal analysis

FC: formal analysis

## Supplementary Information

**SI Fig 1:**
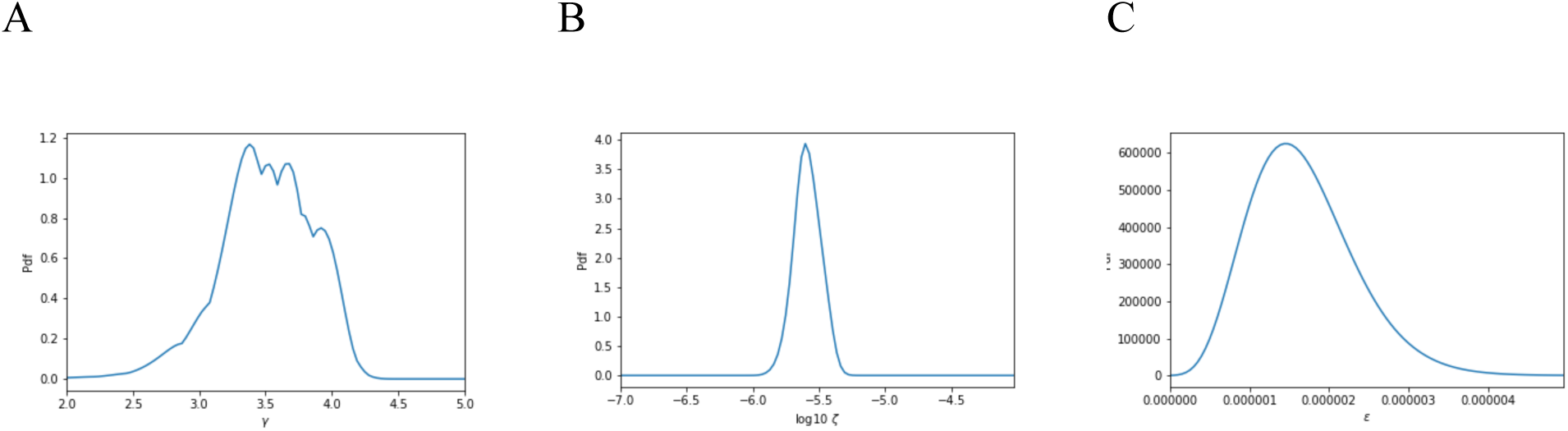
Posterior distributions for the *γ*, ξ, and *ε* parameters for the epidemiological model as shown in Fig 3A (observed site detection rates, for the full archaeological dataset and the combined population estimates in [28,29])

**Supplementary Information Table 1:**
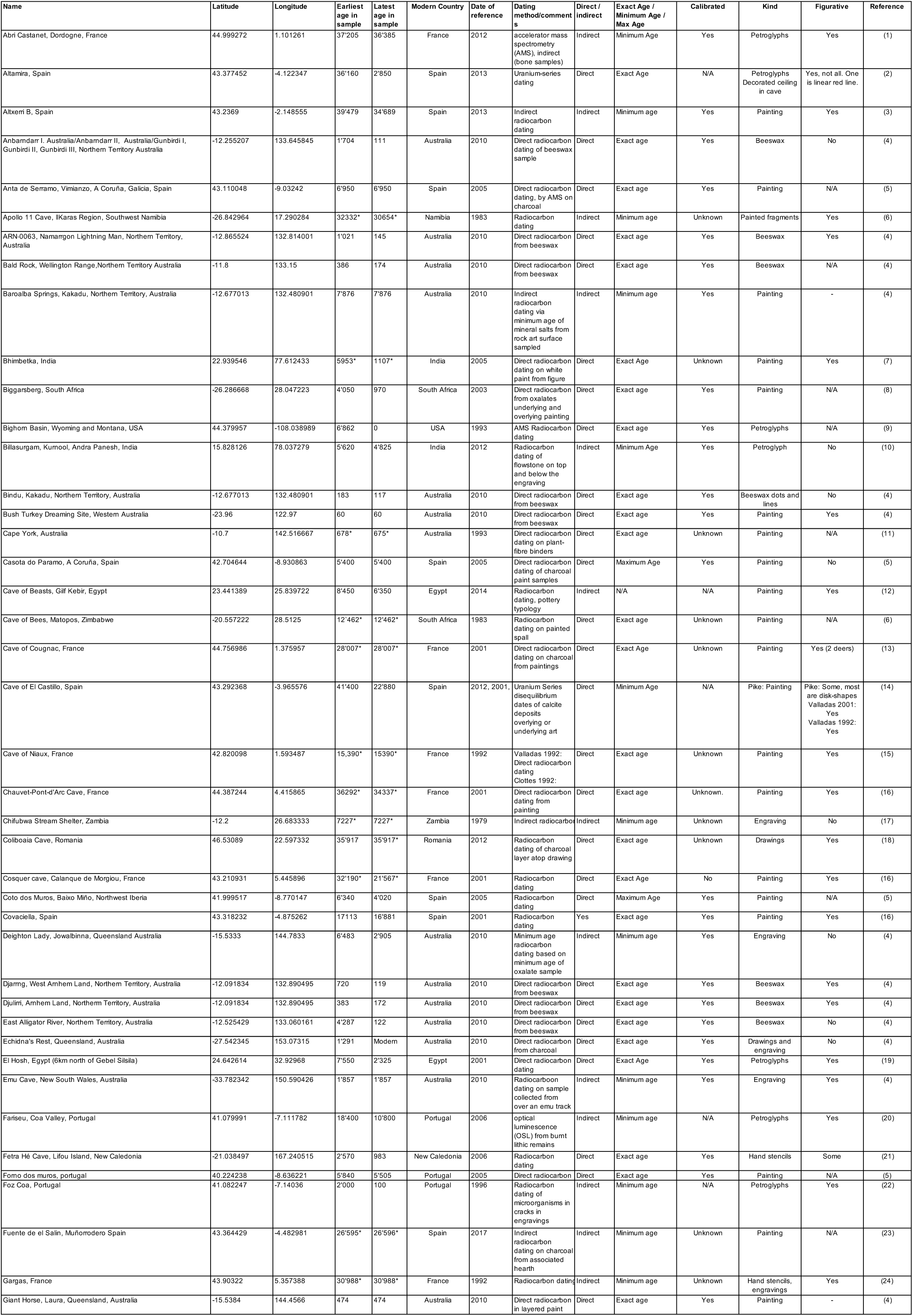
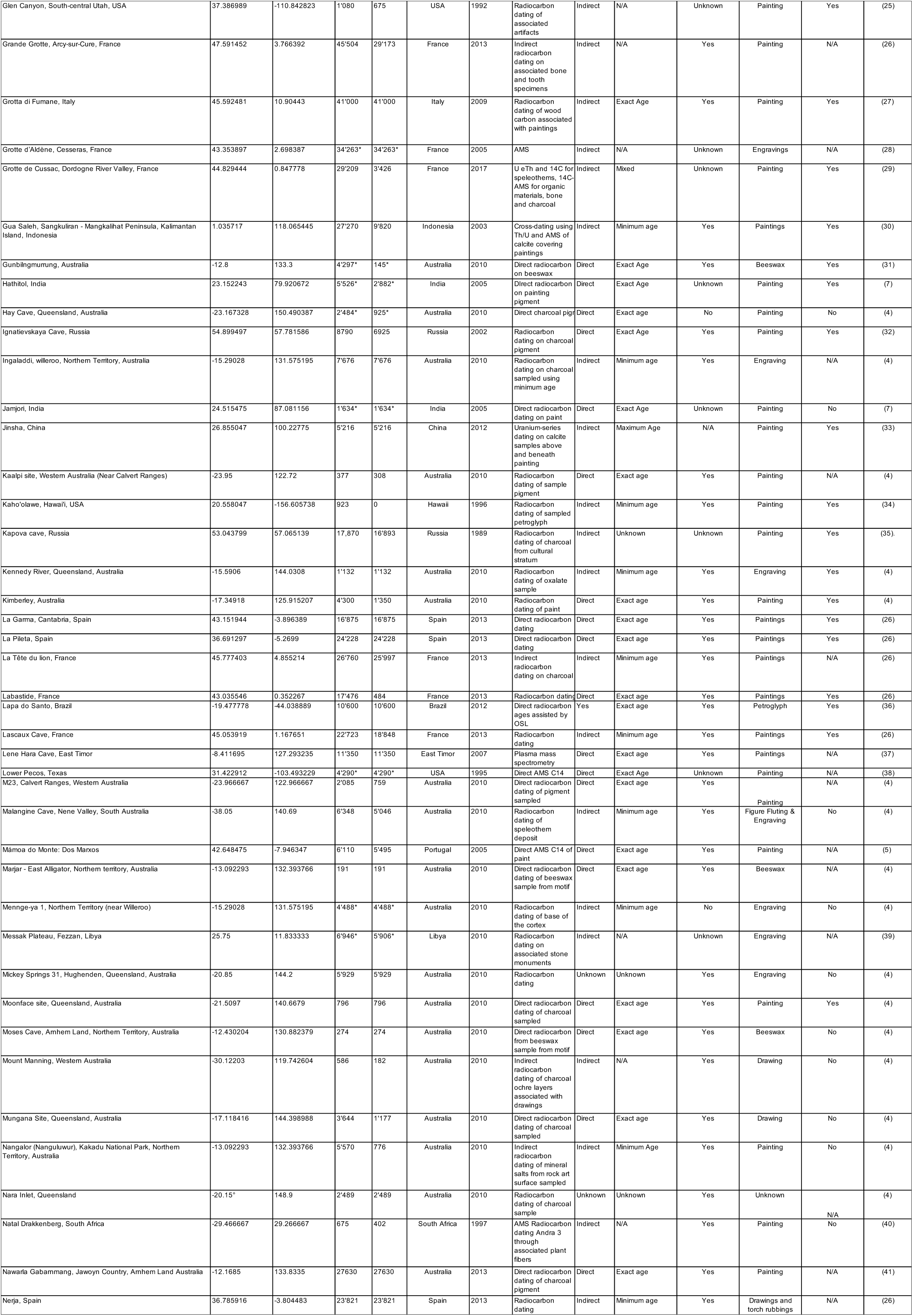
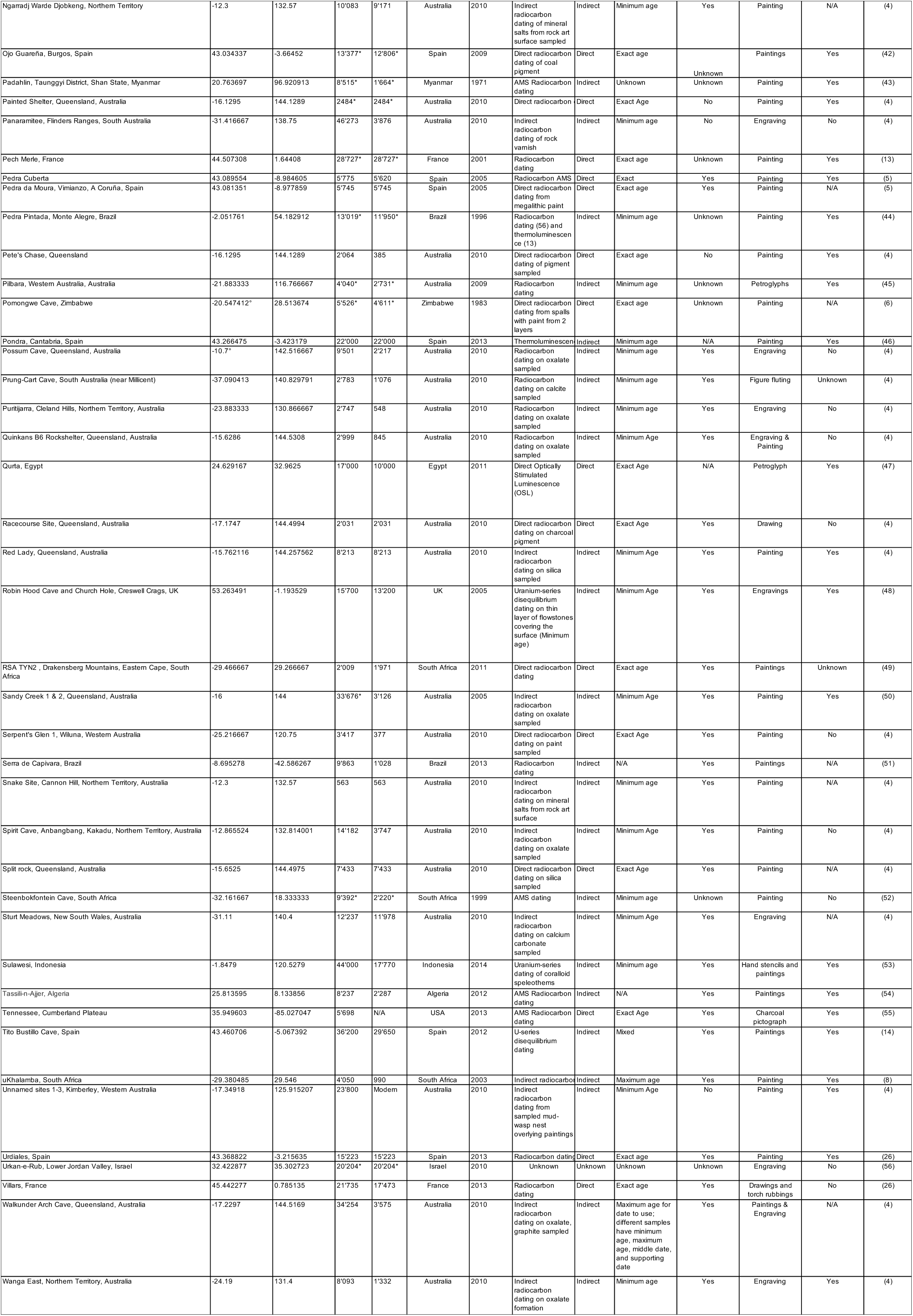
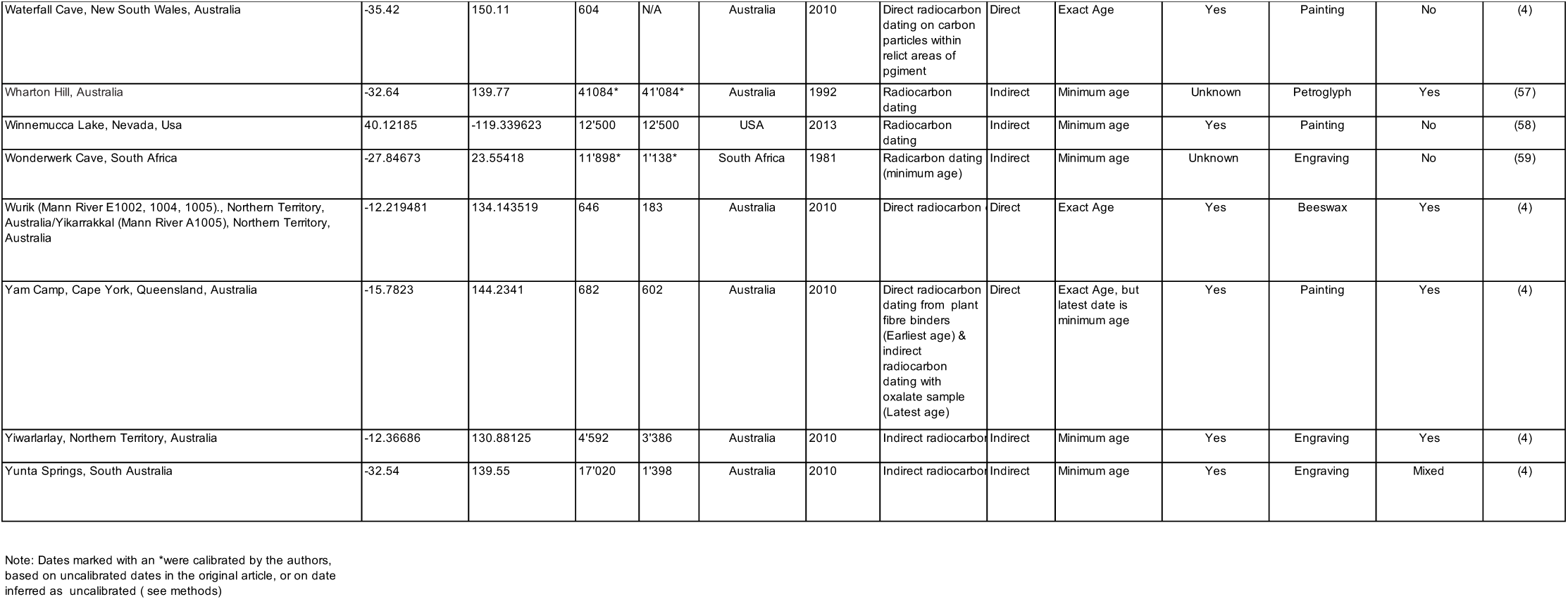
Rock art dataset (133 sites)

